# Comparison of hiPSC-derived hepatic organoids and liver-on-a-chip systems reveal microenvironment-driven maturation

**DOI:** 10.64898/2026.07.28.741157

**Authors:** Isabel Tamargo-Rubio, Tim Krempel, Victoria E.J.M. Palasantzas, Bella Green, Gwen D.L. Weijer, Renée Moerkens, Channen van der Woude, Sven C. D. van Ijzendoorn, Daan J. Touw, Joanne A. Hoogerland, Sebo Withoff, Jingyuan Fu

## Abstract

Human liver organoids (HLOs) and liver-on-a-chip (LoC) systems are emerging as physiologically relevant human models for studying liver function, disease, and drug metabolism, often in combination with human induced pluripotent stem cell (hiPSC)-derived tissues. However, hiPSC-derived models often display batch-to-batch variation and incomplete maturation, and the contribution of microfluidic flow to hepatic maturation remains insufficiently characterized. Here, we developed a cryopreservable and scalable workflow to generate hiPSC-derived hepatic organoids that can be directly matured to either static HLOs or LoC systems, enabling matched comparison of both platforms. Transcriptomic and functional characterization revealed progressive hepatic maturation during organoid differentiation, including increased expression of liver-specific metabolic pathways, enhanced albumin secretion, and increased CYP3A4 activity. Compared to mature HLOs, LoCs exposed to continuous microfluidic flow exhibited transcriptomic profiles suggesting further maturation, with increased enrichment of pathways related to lipid metabolism, xenobiotic metabolism, transport, and tissue organization. These findings demonstrate that microfluidic perfusion promotes hepatic metabolic specialization compared to static organoid culture while maintaining donor-specific characteristics. Together, this study establishes a robust hiPSC-derived LoC platform and highlights the potential of flow-based systems for improved modeling of human liver physiology, disease mechanisms, and drug responses.

## 1. Introduction

Over the past decade, organoids and organ-on-a-chip (OoC) systems have emerged as advanced *in vitro* models with the potential to recapitulate the cellular composition, architecture, microenvironment, and drug sensitivity of *in vivo* organs [1,2]. Among these models, liver-on-a-chip (LoC) models are particularly attractive for biomedical research because access to human liver tissue is limited. Moreover, experiments based on animal models or 2D-cultures of cancer-derived (e.g., HepG2, HepRG) or primary-hepatocyte-derived (e.g., Fa2N4 or IHH) immortalized cell lines often fail to fully recapitulate human liver physiology [3]. Compared to the static culture of organoids, OoC systems introduce dynamic perfusion, spatial control, and multicellular culture under precisely defined microenvironmental conditions, providing a more physiologically relevant alternative. For example, prior work on human hepatocytes cultured under microfluidic conditions showed not only prolonged lifespan and enhanced functionality but also human-specific drug-induced liver toxicities that animal models (such as rat and dog) failed to predict [4,5].

Focus has also shifted towards development of hepatocyte-like cells derived from human induced pluripotent stem cells (hiPSCs). This application of hiPSCs is particularly compelling because they retain the human donor’s genetic background, which is interesting when they are derived from patients with complex genetic diseases. Moreover, hiPSC’s amenability to genetic engineering opens up new opportunities for personalized medicine. Studies have now shown that hiPSC-derived human liver organoids (HLOs) partially recapitulate key hepatic functions, including lipid accumulation, inflammatory signaling, fibrogenic responses, and lysosomal dysfunction [6]. They have also demonstrated their applications in predicting acute hepatoxicity, assessing drug safety, modeling disease phenotypes, and validating therapeutic methods for treating genetic disorders through CRISPR-based gene editing [7–10]. However, hiPSC-derived organoids still exhibit an immature, fetal-like phenotype with reduced metabolic and drug-metabolizing functions [11,12], and a number of studies have aimed to improve hiPSC-derived HLO maturation [10][13–16].

As the complete experimental procedure from hiPSC to LoCs remains highly labor-intensive, costly, and technically sensitive, leading to substantial variability and limited reproducibility, we sought to establish a robust hiPSC-derived LoC platform by further differentiating hiPSC-derived organoids under microfluidic flow conditions to enhance their maturation and functionality. Through parallel generation and matched comparison of hiPSC-derived static organoids and microfluidic flow-based liver-on-chip, we also aimed to improve the reproducibility of hiPSC-based models, explore the opportunities for scalable production, and assess donor-to-donor variability. The resulting hiPSC-derived LoC platform can serve as a personalized *in vitro* model for drug metabolism and disease modeling.

## 2. Results

### 2.1 Generation of expandable and cryopreservable HLOs

To make use of hiPSCs to generate organoid and OoC platforms for biomedical research, we first aimed to establish a reproducible and scalable method to expand liver organoids. Building upon previously published protocols [8,22], we developed and optimized a differentiation procedure to generate expandable HP organoids for three hiPSC lines (two male and one female, Figure 1A−C). The protocol up to the HP stage lasts 23 days and includes three different differentiation stages. In the first stage, DE was successfully induced at day 3, with 70−80% of cells positive for the DE markers CXCR4, GATA4, and SOX17 (Supplementary Figure 1) [23]. In the second stage, cells were differentiated towards HP cells using BMP4 and FGF2 as key growth factors until day 8 (Figure 1A−C) [22,24,25]. In the last stage, organoid GM was added after Matrigel embedding, leading to homogeneous cultures of cystic organoids, which were then cryopreserved for future expansion and culture (Figure 1B, Supplementary Figure 2). This strategy permits scalable pre-production of organoids that can be frozen until needed for further experiments. By generating large batches of HP organoids, technical variability can be reduced.

**Figure 1.**
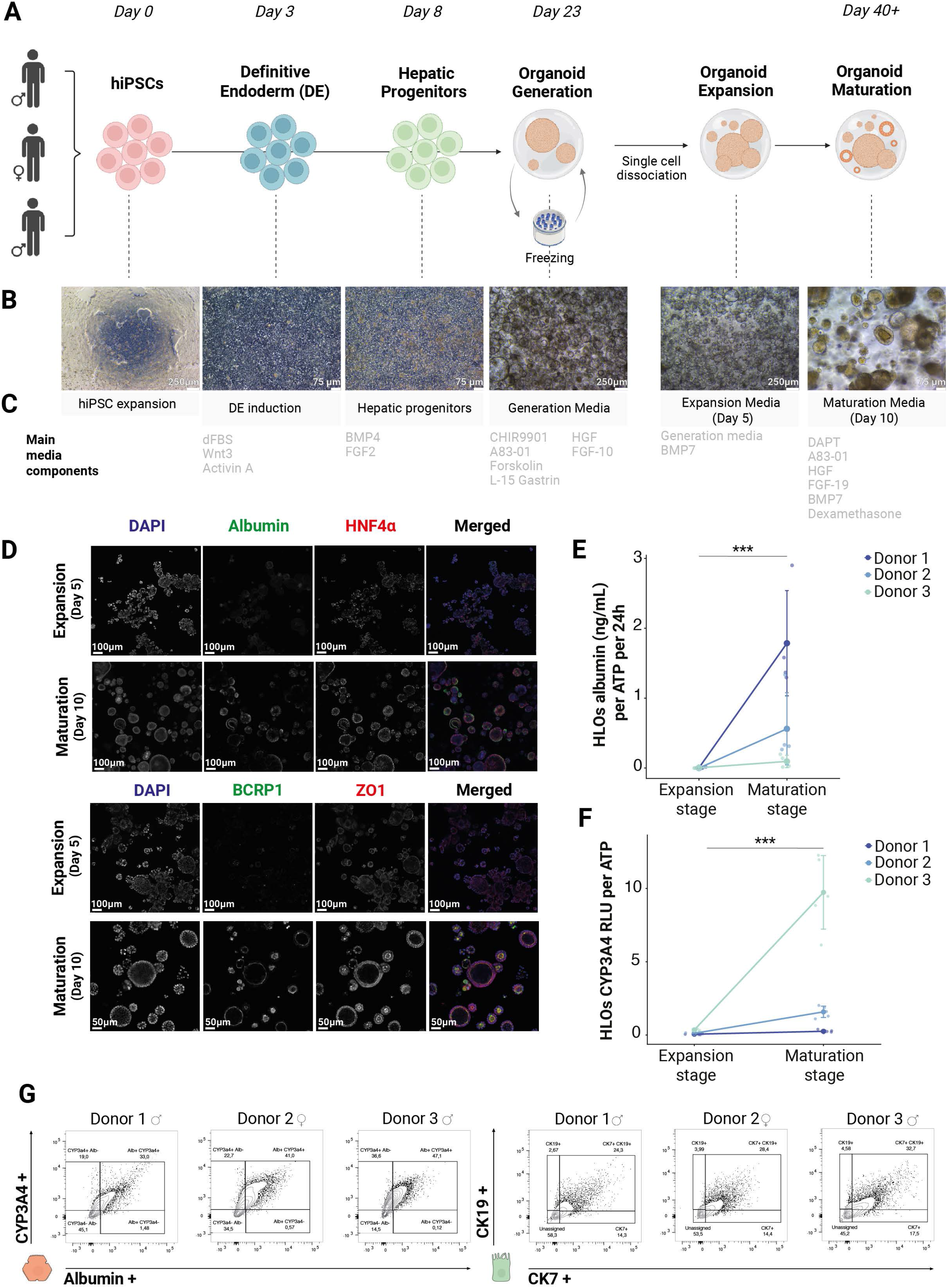
Characterization of human liver organoids (HLOs) di:erentiated from hiPSCs. **(A)** Schematic representation of the di6erent stages of the di6erentiation protocol from hiPSCs to mature HLOs, including a freezing step at the organoid generation stage. **(B)** Representative bright field pictures from Donor 1 of the end of each di6erentiation stage. **(C)** Main media components used at each di6erentiation stage. **(D)** Immunofluorescence staining of hepatocyte markers albumin, BCRP, HNF4α, and ZO1 in HLOs from Donor 1 at the expansion and maturation stage. **(E)** Significant increase (p = 0.00121) in albumin production in 24h between the expansion and the maturation stages in HLOs. Values are corrected for ATP per technical replicate (n = 4 technical per stage and donor). **(F)** Significant increase (p = 0.000275) in CYP3A4 activity between the expansion and maturation stages in HLOs. Values are corrected for ATP per technical replicate (n = 5 technical per stage and donor). **(G)** Flow cytometry analysis for hepatocyte markers CYP3A4 and ALB and for cholangiocyte markers CK7 and CK19. Double positive cells were defined as hepatocyte-like (CYP3A4+, ALB+) or cholangiocyte-like (CK7+, CK19+) cells.

Upon thawing and pre-culturing in GM, organoids can be dissociated and re-seeded as single cells (for speed of growth control) and cultured in EM. In the expansion stage, single cells rapidly develop into organoids within Matrigel domes in 4−5 days while still maintaining their cystic phenotype (Figure 1B, Supplementary Figure 2). Moving into the last stage—the maturation stage—after 10 days in MM, the morphology of most of the organoids changes to form denser, non-proliferative organoids characterized by a clear outer layer of epithelial cells (MM Day 10, Supplementary Figure 2) [7,8]. Although most organoids showed this dense phenotype, some cystic organoids were still present at the end of maturation (red arrows in Supplementary Figure 2). Based on previous literature [26,27] we hypothesize that these hollow, cystic organoids represent progenitor organoids that are more cholangiocyte-like than hepatocyte-like.

### 2.2 Hepatocyte phenotype and cell type composition in mature HLOs

To confirm hepatic differentiation and maturation, we first compared the presence of hepatocyte-like cells at the end of the expansion stage (EM Day 5) and the end of the maturation stage (MM Day 10) (Figure 1D). The number of nuclei positive for the hepatocyte-specific transcription factor HNF4α was low in the expansion stage but markedly increased in the maturation stage. The presence of HNF4α-positive cells in proliferative progenitor organoids is not unexpected as HNF4α also regulates the transition from endoderm to hepatic lineage [28]. Concurrently, albumin-positive cells were absent during expansion but readily detectable after maturation. This was confirmed by albumin protein measurements, where a significant increase was observed between the expansion and maturation stages (p = 0.00121) after correcting for donor effects (Figure 1E). Mature organoids also displayed a polarized phenotype, as evidenced by the high density of apical BCRP1 in the inner side (Figure 1D). We further confirmed that CYP3A4 enzyme activity, a major hepatocyte marker for xenobiotic metabolism, is significantly enhanced (p = 0.000275) at the maturation stage (Figure 1F). Together these results demonstrate hepatic specification and maturation, with enhanced hepatocyte-specific metabolic activity.

We investigated our system for the presence of other cell types. Flow cytometry revealed a 33−47% ALB^+^CYP3A4^+^-positive hepatocyte-like population and a 5−10% CK7^+^CK19^+^-positive cholangiocyte-like population (Figure 1G). We also detected small populations of liver sinusoidal endothelial cells (CD14^+^LYVE1^+^, 0.1−0.3%) and stellate cells (CD56^+^, αSMA^+^, 0.2−0.6%), ranging from 0−0.6% across different donors. Kupffer cells (CD45^+^, CD68^+^, MHCII^+^) categorized under the term ‘immune cells’ were not found. (Supplementary Figure 3, Supplementary Table IV).

While we observed high reproducibility among technical replicates from the same donor, the variability between donors was notable. Donor 1 showed the highest albumin production, while Donor 3 showed the highest CYP3A4 activities (Figure 1E and F). In terms of cellular proportions, Donor 3 displayed the highest percentage of hepatocyte-like cells and the lowest percentage of cholangiocyte-like cells, whereas Donor 1 showed the opposite pattern.

### 2.3 Enhanced liver-specific molecular pathways in mature HLOs

To further assess HLO maturation, we carried out comparative transcriptomic analysis for the expansion (EM Day 5) and maturation stages (MM Day 10). PCA revealed a clear separation between expansion and maturation stage samples. PC1 explained the largest proportion of variance and primarily distinguished the two differentiation stages (Supplementary Figure 4A), whereas PC2 captured the variability between donors (Supplementary Figure 4A). Differential gene expression analysis identified 1992 DEGs, with the albumin-encoding *ALB*-gene emerging as one of the most strongly upregulated genes in the maturation stage (Supplementary Figure 4B).

Differential expression analysis revealed two distinct sets of DEGs: genes upregulated after the expansion stage and genes upregulated at the end of the maturation stage (Figure 2A-2D). Pathway analysis on genes upregulated after the expansion stage showed enrichment for pathways related to cell division and proliferation, such as DNA replication, cell cycle checkpoints, chromosome segregation, and rRNA processing (Figure 2B). Among these DEGs, the most biologically relevant included proliferation markers (e.g., *MKI67*), master cell-cycle regulators (e.g., *AURKA*, *CCNA2*, *PLK1*), genes related to mitotic checkpoint regulation (e.g., *CDC20*, *BUB1*), and genes related to chromosome topology (e.g., *TOP2A*) and segregation (e.g., *SGO1*) (Figure 2D). The gene set upregulated at the end of the maturation stage was enriched for pathways related to key hepatic functions, including lipid homeostasis, transport and metabolism; steroid and xenobiotic metabolism; coagulation and hemostasis; and extracellular matrix (ECM) and development (Figure 2C). Among these DEGs, notable genes included lipid and cholesterol transporters (e.g., *ABCA5*, *APOB*), the bile acid transporter *SLC10A1*, and lipid metabolism genes (*APOA1* and *APOE*) (Figure 2D), together with genes involved in xenobiotic response, steroid and xenobiotic metabolism, and cellular glucuronidation.

**Figure 2.**
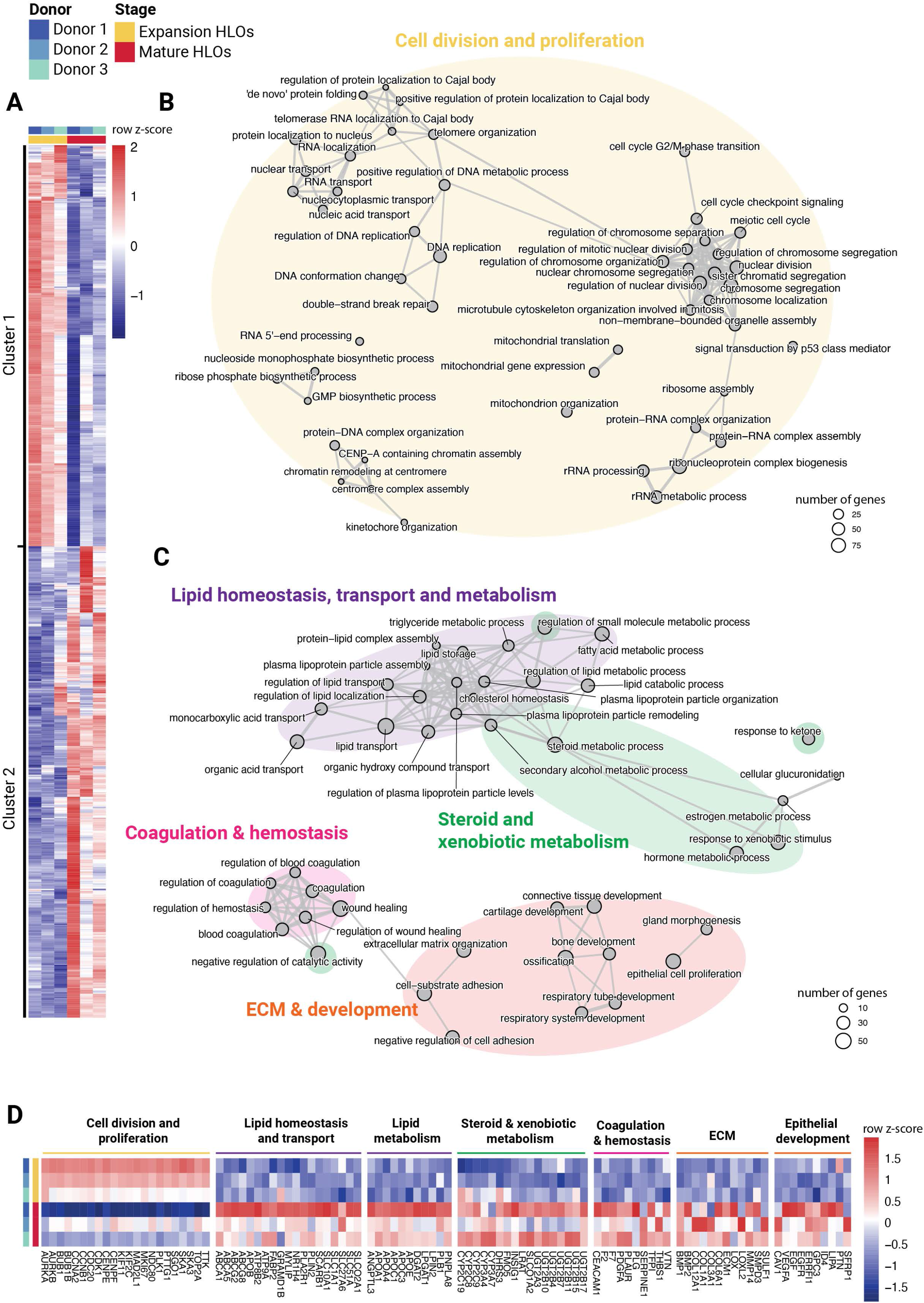
Multi-pathway transcriptomic analysis of HLOs at the end of the expansion and maturation stages. **(A)** Heatmap of average expression of di6erentially express genes (DEGs) between Expansion HLOs (yellow) and Mature HLOs (red), clustered by expression profile. Cluster 1 contains genes upregulated in Expansion HLOs. Cluster 2 contains genes upregulated in Mature HLOs. Color indicates row z score. Columns correspond to donors (left = Donor 1, middle = Donor 2, right = Donor 3). Technical replicates n = 6 donor and stage. **(B)** Network of enriched biological processes for the DEGs upregulated in Cluster 1 (upregulated in Expansion HLOs). **(C)** Network of enriched biological processes for the DEGs upregulated in Cluster 2 (upregulated in Mature HLOs). ECM; extracellular matrix. In (B) and (C), the top 50 biological processes with lowest adjusted *p* value that belong to the cluster are shown. Node size indicates the number of DEGs in each pathway. Thickness of the lines connecting the nodes represents the number of shared DEGs. **(D)** Heatmap of average expression of DEGs selected from the enriched biological processes described. Color represents column z score. Rows correspond to donors (top = Donor 1, middle = Donor 2, bottom = Donor = 3) for each of the two conditions (see color key at top left).

Upon maturation, we observed a marked induction of major phase I CYP450 enzymes and phase II UDP-glucosyltransferases family members. In line with increased CYP3A4 enzyme activities (Figure 1F), CYP3A4 gene expression showed an 18.9-fold increase in mature HLOs compared to the expansion stage. Notably, the donor-specific enzyme activity differences we had observed were also reflected at expression level (Figure 2D), with Donor 3 exhibiting the highest expression levels and Donor 1 the lowest.

In addition to pathways related to hepatic function, we also observed an enrichment in genes involved in coagulation and hemostasis (e.g., key blood coagulation factors *F2*, *F7*, and *TFPI*) which also reflect a more mature hepatocyte phenotype [29], epithelial development and ECM organization (e.g., *COL1A1*, *COL3A1*, and *COL6A1*), and genes involved in developmental signaling and epithelial or progenitor cell differentiation and maturation (e.g., *GPC3*, *ID4*, and *SFRP1*) (Figure 2D) [30,31].

Altogether, these results suggest that the organoids may still exhibit active multicellular remodeling and a developmental-like state, driven by a mixture of hepatocyte-like, progenitor-like, and non-parenchymal cells The gene signatures we observed likely reflect ongoing tissue differentiation, organization and stromal/epithelial interactions.

### 2.4 Hepatic maturation under continuous and unidirectional microfluidic flow

Previous work in our lab had established that hiPSC-derived intestinal epithelial cells differentiate further on microfluidic chip than when seeded in static culture vessels in Matrigel domes or in Transwell systems [17]. As we had observed a rather incomplete maturation stage (as shown above), we decided to test the hypothesis that hepatocytes will also differentiate further under microfluidic flow conditions on chip. For this, cryopreserved HP organoids were thawed, expanded, and dissociated into single cells and seeded into the top channel of Emulate S1 chips. Cells were cultured on chip for approximately 20 days (5 days in EM and 15 days in MM) using the same media as before (Figure 3A-3C). On-chip, the cells evolved from a thin monolayer at the end of the expansion stage to a thicker layer at the end of the maturation stage (Figure 3B), forming 3D structures in some areas of the channels (Figure 3B black arrows, Supplementary Figure 5). The mature phenotype became apparent between days 5−7 of the maturation stage and persisted throughout, allowing cells to survive up to 14 days (Figure 3B, Supplementary Figure 5). Albumin production increased significantly (p = 0.000339) at the end of the maturation stage relative to the expansion stage, and we observed donor-to-donor variation comparable to that seen in HLOs (Figure 3D, Supplementary Figure 4C). CYP3A4 activity was also detected in the chips for all three donors (Supplementary Figure 6A), but further validation using CYP3A4-specific drugs will be necessary to confirm these results.

**Figure 3.**
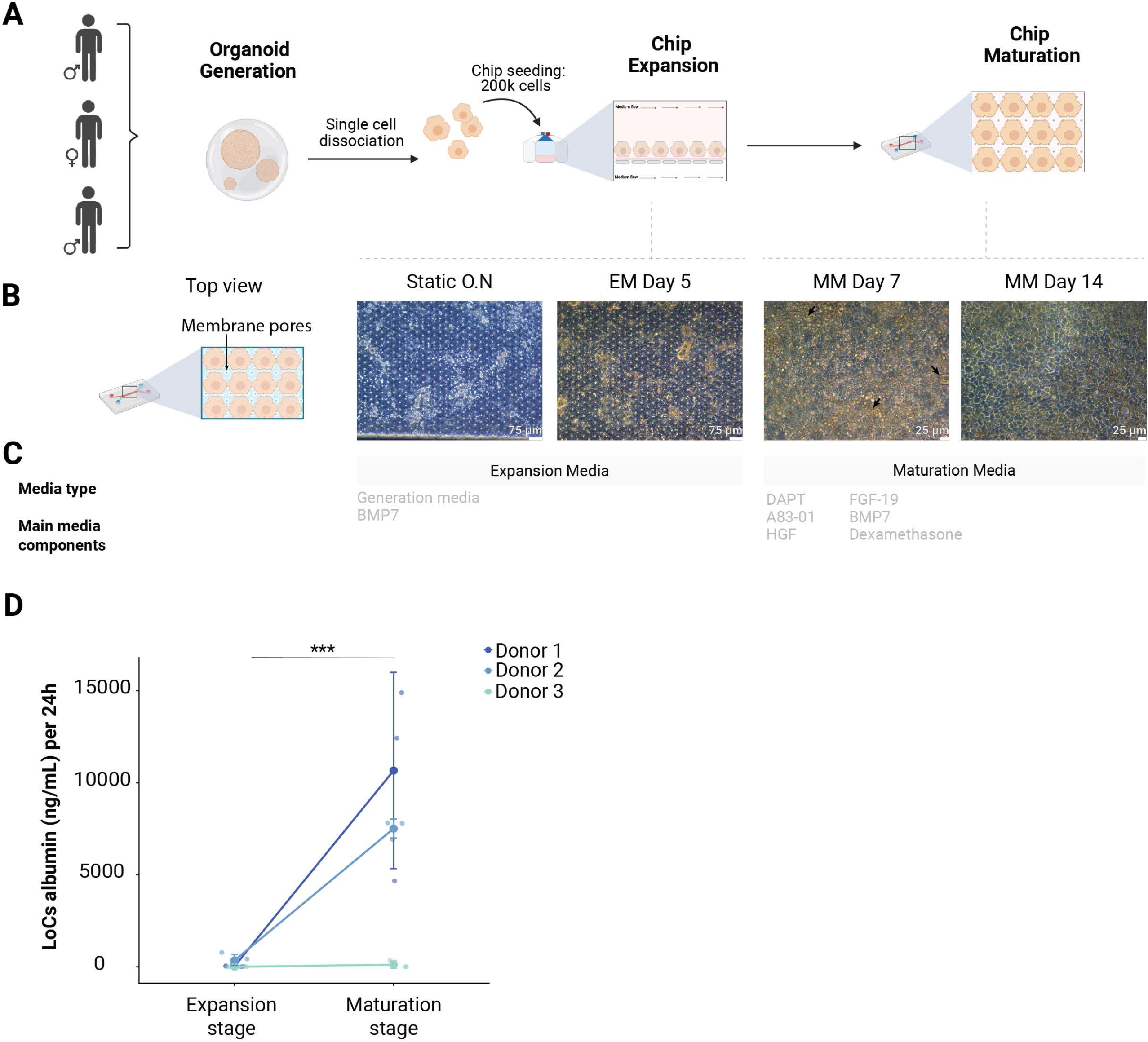
Differentiation protocol from expandable HLOs to mature liver-on-a-chip (LoC). **(A)** Schematic representation of the di6erent stages of the di6erentiation protocol from hiPSCs to mature LoCs. **(B)** Representative bright field images (top view) from LoC (Donor 1) throughout the expansion and maturation stages. At maturation media (MM) Day 7, some 3D structures start appearing (black arrows). **(C)** Main media components used at each di6erentiation stage. **(D)** Significant increase (p = 0.000339) in albumin production in 24h between the expansion and the maturation stages in LoCs. Media collected from the same LoC over time. Technical replicates in expansion media n = 4. Technical replicates in maturation media n = 3.

### 2.5 Enhanced metabolic specialization in LoC

We compared the transcriptional profiles of mature LoCs with those of mature HLOs grown under static conditions. Here we observed a pronounced donor effect in the LoCs, with PC1 from the PCA clearly separating the samples from different donors (Supplementary Figure 6B). After including PC1 as a covariate in subsequent analyses, a DEG analysis comparing mature HLOs to mature LoCs indicated enhanced maturation in the LoC cultures (Figure 4A-4D).

**Figure 4.**
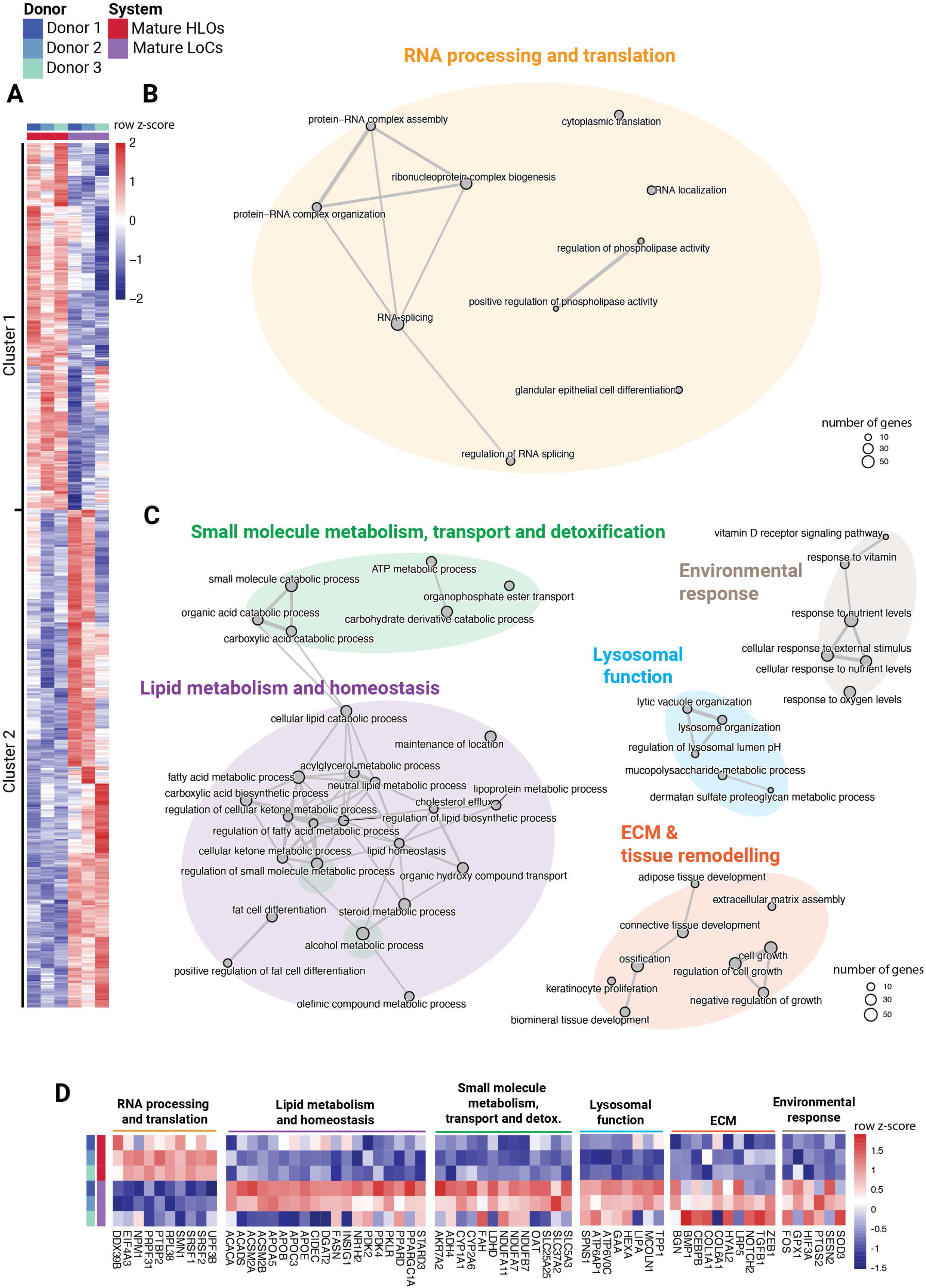
Multi-pathway transcriptomic analysis of LoC and HLOs at the end of the maturation stage. **(A)** Heatmap of average expression of DEGs between Mature HLOs (red) and Mature LoC (purple), clustered by expression profile. Cluster 1 includes genes upregulated in Mature HLOs. Cluster 2 contains genes upregulated in LoCs. Color indicates the row’s z score. Columns correspond to donors (left = Donor 1, middle = Donor 2, right = Donor 3). Technical replicates: n = 6 for Mature HLOs and n = 3 for Mature LoCs. **(B)** Network of enriched biological processes for the DEGs upregulated in Cluster 1 (upregulated in Mature HLOs). **(C)** Network of enriched biological processes for the DEGs upregulated in Cluster 2 (upregulated in LoC). ECM; extracellular matrix. In (B) and (C), a maximum of 50 biological processes with the lowest adjusted *p* value that belong to the cluster are shown. **(D)** Heatmap of average expression of DEGs selected from enriched biological processes. Color represents column z score. Rows correspond to the individual donors (top = Donor 1, middle = Donor 2, bottom = Donor 3) for each of the two models (see color key at top left).

In LoCs, downregulated genes were involved in RNA processing and post-transcriptional regulation, such as RNA splicing and protein-RNA complex assembly and organization (Figure 4B). Example genes included *SRSF1*, *NPM1,* and *RLP38* (Figure 4D). This indicates higher transcriptional activity in HLOs compared to LoCs, suggesting ongoing differentiation programs and less terminal maturation in HLOs. By contrast, the genes upregulated on chip (Figure 4C) pointed to more mature liver functionality. These included genes involved in lipid metabolism and homeostasis, small molecule metabolism, transport and detoxification, lysosomal function, ECM, tissue remodeling and development, and environmental response (Figure 4B). In particular, genes involved in phase I and phase II metabolism (e.g., *CYP1A1*, *CYP2A6*, *ADH4*) were enriched in LoCs compared to HLOs. This was accompanied by increased expression of additional detoxification-related genes, indicating an overall upregulation of xenobiotic and aldehyde metabolism pathways. These findings are consistent with enhanced hepatocyte functional capacity under perfused conditions [5]. In addition, genes involved in lipid metabolism were prominently enriched in LoCs, including key regulators of lipid synthesis and storage (e.g., *FASN*, *DGAT2*, *APOB*). Notably, *APOB*, which encodes the structural protein essential for very low-density lipoprotein assembly and secretion, was upregulated. Lipoprotein assembly and export are central hepatic functions that are often poorly recapitulated in static *in vitro* liver models [32] because physiological lipoprotein metabolism is challenging to maintain outside a dynamic microenvironment.

Together, our findings indicate that microfluidic perfusion enhances metabolic maturation and tissue-level organization, improving lipid handling, xenobiotic metabolism, and additional metabolic processes.

## 3. Discussion and Conclusion

Here, we describe a robust, time-efficient method for generating hiPSC-derived HLOs and LoC platforms with strong potential for human-based biomedical research. Although the complete differentiation protocol requires ∼40 days, our incorporation of a validated (optional) cryopreservation step enables a freeze–thaw workflow for scalable organoid production. This shortens the subsequent differentiation period upon thawing to only ∼15–20 days and reduces technical variation by generating big batches of HP organoids that can be cryopreserved.

When comparing the end of the expansion and maturation stages, the hiPSC-derived HLOs we generated presented a transcriptomic shift from more proliferative progenitor programs to functional hepatocyte programs, recapitulating key liver functions involved in phase I and phase II drug metabolism, lipid/bile transport, and metabolism. These features, which define the core detoxification and metabolism machinery of hepatocytes, have been recognized as hallmarks of hepatocyte maturation [33–35]. We also identified enrichment of pathways related to coagulation and complement cascade in our mature HLOs. The proteins associated with these pathways are primarily produced by primary hepatocytes and contribute to key biological processes, including immune regulation, liver regeneration, and maintenance of tissue homeostasis [36]. Co-enrichment of these functions, which are interdependent *in vivo*, is unlikely to occur without real hepatic specification, indicating successful hepatic maturation.

Organoids have the potential to recapitulate the cellular composition and architecture of the liver. Our HLO cell populations were dominated by hepatocyte-like cells (33–47% ALB⁺CYP3A4⁺). When including cells positive for at least one of these hepatocyte-defining markers (albumin and CYP3A4), this proportion increases to 53–83%, close to the 60% hepatocytes reported for the liver [31]. The presence of mature hepatocytes is also supported by increasing albumin secretion and CYP3A4 activity. In addition to hepatocytes, 5−10% of the cell population in our HLOs belonged to cholangiocytes (5–10% CK7⁺/CK19⁺ cells). The presence of both cell types is consistent with the fact that cholangiocytes and hepatocytes both originate from a common progenitor. Our findings are also consistent with previous reports showing that hiPSC-derived liver organoids frequently contain both epithelial liver lineages, which are organized into hepatocyte-like regions and bile duct–like structures formed by cholangiocytes [8,37]. Moreover, we also observed a very small proportion of other liver-resident cell types, including liver sinusoidal endothelial cells and Kupffer cells. The coexistence of multiple cell types highlights the cellular complexity and physiological relevance of our model.

Compared to the HLOs, our LoC platform exhibited further enhanced hepatic functions under microfluidic flow, in agreement with previous findings [17,38]. For example, we observed downregulation of RNA processing and translation programs, indicating that the cells on chip exited a plastic state and progressed towards a terminally differentiated state. The chip environment also appears to generate more comprehensive lipid handling, transport, and metabolism functions. Upregulated genes indicated enhanced small molecule metabolism, coupled to increased bioenergetic capacity through oxidative phosphorylation, consistent with previous findings in hiPSC-derived hepatocyte-like cells cultured on chip from Danoy et al. [39]. These are all characteristics of improved hepatic maturation on chip.

Maturation in LoC was also evidenced by upregulation of genes related to oxidative phosphorylation [40], nutrient and oxygen sensing, and environmental responses. Similar pathways were also found enriched in an hiPSC-derived gut-on-chip compared to static organoid culture [17] and in kidney organoids cultured under flow conditions [38]. These consistent observations across different organ systems suggest that continuous media perfusion under microfluidic flow conditions prevents nutrient depletion and waste accumulation, thereby promoting the nutrient-sensing and environmental response pathways that are essential for cellular function and maturation. Notably, while flow-mediated improvements compared to static conditions have generally been modest across different organ models [41], highly perfused organs like the liver show stronger responses, with these changes apparent when checking certain liver-specific markers (e.g., CYP3A4 expression and BSEP) [41]. In line with these findings, our broader multi-pathway transcriptomic analysis demonstrates enhanced maturation in LoCs that extends beyond the isolated biomarker improvements typically reported in literature [42–46].

Donor-to-donor variability in hiPSC generation and differentiation is one of the important technical considerations in hiPSC-based biological models. In our datasets, we also noted significant inter-donor variation, which explained 26.6% of the transcriptional variation in organoids and 48.9% of the variance in LoCs. This variation could be due to donor differences in genetic make-up, epigenetic memory, and clonal variation. A previous study reported that approximately 50% of the genome-wide expression variability in their 317 hiPSC lines was driven by genetic background [47], and the authors showed that this donor-specific transcriptomic profile was preserved through both reprogramming and hepatic differentiation. Although the origin cell type for iPSC-generation can influence transcriptomic profiles [48], donor-dependent effects generally outweigh the variability introduced by the starting cell type or reprogramming method [49]. Previous studies using standardized hepatic differentiation protocols across 24 hiPSC lines similarly demonstrated reproducible within-line differentiation while retaining inter-individual differences in CYP activity characteristic of primary hepatocytes [50].

Another potential contributor to inter-donor differences is epigenetic variation, which becomes more apparent upon differentiation [51]. Supporting this, Park et al. demonstrated that CpG hypermethylation and histone modifications restrict CYP450 expression in hepatocytes derived from human embryonic stem cells and that combined inhibition of DNA methyltransferases and histone deacetylases partially restored CYP1A1 and CYP1A2 expression [52]. Consistent with these findings, we observe donor-dependent differences in albumin secretion and CYP3A4 activity in both HLOs and LoCs. Together, these results indicate that a combination of genetic and epigenetic determinants may explain the inter-donor variability we observe in our systems. The maintenance of these donor-specific characteristics highlights the potential of this platform for personalized medicine applications, including pharmacogenomics and disease modeling, and underscores the importance of including multiple donors during model development. Expanding the donor panel to include clinically relevant pharmacogenomic variants, such as CYP2D6 poor or extensive metabolizers, could help determine whether chip-mediated maturation effects are consistent across genetic backgrounds under flow conditions.

Beyond promoting hepatic maturation, LoC architecture enables experimental approaches and extends culture longevity beyond that of static organoid systems (10 days for mature HLOs vs 15 days for mature LoCs in this study). Spatially separated channels allow for integration of additional cell types in complex co-cultures with independent media [53,54], enabling incorporation of physiologically relevant liver compartments [38,55]. Addition of non-parenchymal cells, including Kupffer, stellate, and sinusoidal endothelial cells, from the same hiPSC donor can introduce immune, fibrogenic, and vascular components for advanced disease modeling. Continuous flow also enables longitudinal sampling and removal of toxic metabolites, supporting studies of chronic drug exposure and diseases such as metabolic dysfunction-associated steatotic liver disease. Combining OoC and hiPSC technologies enables modeling of inter-organ communication using donor-matched organ chips, thereby preserving the donor’s genetic background while improving the study of physiology, disease progression, and pharmacokinetic/pharmacodynamic responses.

In conclusion, our method enables the generation of cryopreserved HP organoids from hiPSCs, and these organoids can be further expanded and matured to produce mature HLOs and LoCs. We further show highly reproducible results between different technical replicates within the same donors, while donor-to-donor variation is significant. Hepatic maturation and functionality were confirmed in our models through transcriptional analysis and functional assays. Despite both platforms showing biologically relevant hepatic function, LoCs showed a more complete metabolic transcriptomic profile than HLOs, indicating that chip platforms might be more suitable for recapitulating hepatic function (especially lipid metabolism). However, these results need to be further validated and expanded with more functional assays adapted to chip systems. Together, our findings support the growing role of HLOs and LoC systems as next-generation models for human liver research.

## 4. Materials and Methods

### 4.1 hiPSC reprogramming and culturing conditions

Three hiPSC lines were generated as previously described by Moerkens et al. [17]. hiPSC lines were maintained in mTeSR™ Plus (Stemcell Technologies, #05825) on 6-well plates coated with hESC-qualified Matrigel (Corning, #354277). Cells were passaged every 3−4 days using ReLeSR (Stemcell Technologies, #05872), according to the manufacturer’s instructions, and cultured in a humidified environment at 37°C in 5% CO_2_. hiPSC lines were then cryopreserved in CryoStor CS10 (Stemcell Technologies #7930). All experiments with hiPSC lines were approved by document no. METC 2013/440.

### 4.2 Hepatic progenitor differentiation and organoid generation

Before differentiation, hiPSC lines were thawed and cultured in mTeSR™ Plus (Stemcell Technologies, #100-0276) on Nunc™ Cell Culture−treated 6-well plates (Thermo Fisher Scientific, #140675) at 37°C in 5% CO_2_. When 70% confluency was reached, cells were dissociated with StemPro® Accutase® Cell Dissociation Reagent (Thermo Fisher Scientific, #A1110501), seeded as single cells at a density of 5x10^4^ cells per well on Nunc™ Cell Culture−treated 24-well (Thermo Fisher Scientific, #142475) Matrigel® coated plates, and cultured in mTeSR™ Plus. When 50−80% confluency was reached (optimized per donor), cells were differentiated toward definitive endoderm (DE) by changing the media each day for 3 consecutive days. Day 1 medium consisted of RPMI-1640 containing L-glutamine, Pen/Strep, activin A, and Wnt3a. For culturing days 2 and 3, Wnt3a was omitted and defined fetal bovine serum (dFBS) was added. For subsequent hepatic progenitor (HP) induction, DE cells were cultured in HP media consisting of RPMI-1640 containing B27 supplement, BMP4, FGF2, and DMSO. This media was changed every 24 hours for 5 days.

HP cells were detached by a 6-minute incubation with StemPro® Accutase® Cell Dissociation Reagent and embedded in 10 μL Matrigel® domes (3x10^6^ cells per mL, 3 domes per well) in Nunc™ Cell Culture−treated 24-well plates. After incubation for 10 minutes for Matrigel gelation, generation medium (GM) supplemented with RevitaCell™ was added for 24 hours. GM consists of Advanced DMEM/F12 supplemented with GlutaMAX, HEPES, N_2_, B27, bovine serum albumin (BSA), N-acetyl-L-cysteine, [Leu15]-gastrin-I-human, nicotinamide, A83–01, forskolin, CHIR99021, FGF10, and HGF. This medium was replenished every 2 days for 13–15 days. hiPSC-HLOs were either passaged (1:3) in fragments for serial expansion or frozen until further use. For fragment passaging, pipette tips were coated in 2% FBS in PBS and organoids were released from Matrigel domes by mechanical dislodgement and repeated pipetting in cold Advanced DMEM/F12. After centrifugation (400 × g, 5 minutes, 4^◦^C), the Matrigel was removed and organoids were fragmented by repeated pipetting. After fragment dissociation, HLOs were either re-seeded in Matrigel domes and cultured in GM or frozen in CryoStor (Sigma Aldrich, #C3124).

Details of the media compositions for DE induction, HP induction, and GM, including components, concentrations, product references, and culture times, are described in Supplementary Table I.

### 4.3 Single-cell dissociation, organoid expansion, and maturation

To normalize the amount of starting material after expanding HLOs to a sufficient amount for our experimental conditions, HLOs were dissociated into single cells and re-seeded at the desired density. HLOs in GM were recovered from the Matrigel domes and dissociated in fragments as described above. Fragmented organoids were centrifuged (400 × g, 5 minutes, 4^◦^C) and incubated for 10 minutes in 1 mL of TrypLE (Thermo Fisher Scientific, #12604013) per 12 Matrigel domes. The suspension was carefully resuspended and filtered through a 70 µM filter to remove cell aggregates. Cells were counted and seeded in 10 μL Matrigel domes at a density of 5x10^6^ cells per mL and incubated with expansion medium (EM) comprising GM supplemented with BMP7 for 5 days, changing media every other day. The EM was supplemented with RevitaCell™ for the first 24 hours. For hepatocyte maturation, hiPSC-HLOs were cultured in maturation media (MM) containing Advanced DMEM/F12 supplemented with GlutaMAX, HEPES, B27, BSA, N-acetyl-L-cysteine, [Leu15]-gastrin I human, DAPT, A83–01, dexamethasone, FGF19, BMP7, and HGF for 10 days, changing media every other day. An overview of the full differentiation protocol to HLOs is shown in Figure 1A.

### 4.4 LoC culturing: chip activation and coating, cell seeding, expansion, and maturation

Emulate Organ-Chip was activated using Emulate’s proprietary reagents, 1 mg/mL ER1™ and ER2™, according to the manufacturer’s instructions. Both microfluidic channels were coated with hESC-qualified Matrigel (Corning, #354277). Single-cell dissociated organoids were seeded at a density of 2x10^5^ cells (200K cells/chip) in the top channel. After 2−6 hours of attachment under static conditions in EM media supplemented with RevitaCell™, the chips were washed by gravitational force with the EM media and left static overnight to ensure proper attachment of cells and sufficient growth factor concentrations in the media. The following day, cell attachment was confirmed and chips were washed by gravitational force using degassed EM to remove dead cells before connecting to the flow. Chips were then attached to the Pod™ Portable Module and placed in the Zoë™ Culture Module. They underwent a regulate cycle to prevent formation of bubbles by increasing the pressure for 2 hours. Fresh degassed EM media was then applied for 4 days. For subsequent cell maturation, the media reservoirs were emptied and degassed MM media was added and replenished every 3 days for 14 days. For media degasification, culture media was kept at 37°C for at least 1 hour and filtered under vacuum using a Millipore® Steriflip® (Merck, #SCGP00525). Throughout the experiment, we applied a flow rate of 30 µl/h in the top channel and of 10 µl/h in the bottom channel.

### 4.5 Flow cytometry

At the end of DE induction, 1 million dissociated cells for each donor were used for flow cytometry analysis to assess DE purity. A fraction of the cell suspension of the biological replicates in each condition was pooled for use as an ‘unstained sample’ in flow cytometry analysis. Cells were incubated with Accutase for 6−8 minutes. The single-cell suspension was washed and incubated in 100 μL of FcBlock (BioLegend, #422302) for 30 minutes at room temperature (RT). Extracellular staining was performed by incubating the cells with CXCR4-APC for 40 minutes at 4°C. After fixation with 4% PFA, cells were permeabilized and labeled with fluorophore-conjugated intracellular antibody mix containing GATA4-A488 and SOX17-PE in BD Perm/Wash buffer (BD Biosciences, #554723) for 30 minutes at 4°C. Cells were then resuspended in DPBS with FBS (2% v/v) and measured using the Quanteon analyzer at the Flow Cytometry Facility of the University Medical Center Groningen (UMCG).

HLOs were harvested at the end of their maturation stage for flow cytometry analysis following the same single-cell dissociation protocol described in “Single-cell dissociation, organoid expansion, and maturation”. Before proceeding with the antibody staining, cells were washed with cold PBS (300 x g, 5 min, 4°C) and stained with zombie aqua for 30 minutes at 4°C. After incubation, cells were carefully washed with PBS and fixated in 4% PFA for 10 minutes at RT. Single stains (with UltraComp eBeads™ Plus Compensation Beads, Thermo Fisher Scientific, #01-3333-41), fluorescence minus one (FMO) controls, and stained and unstained samples were prepared. Cells were incubated with FcBlock for 30 minutes and then incubated for 40 minutes at 4°C with extracellular staining mix (CD56-BUV395, HLA-DQ-BUV737, CD14-BV650, CD45-BV711, and LYVE1-AF594 diluted in DPBS with FBS (2% v/v) containing BD Horizon™ Brilliant Stain Buffer (BD Horizon, #563794)). After incubation and washing with DPBS with FBS (2% v/v), samples were permeabilized in 1x Perm/Wash buffer for 15 minutes at RT. Samples were then incubated for 30 minutes at 4°C with intracellular antibody mix containing CK7-BV421, CYP3A4-FITC, ACTA2-PerCp, CD68-PE-Cy7, Albumin-PE, CK19-APC, and HNF4α-AF750 in Perm/Wash buffer. After incubation, samples were washed and resuspended in 400 μL of DPBS with FBS (2% v/v) and measured using the BD FACSymphony™ SORP analyzer at the UMCG Flow Cytometry Facility. All washing steps were performed at least twice in cold DPBS with FBS (2% v/v) and centrifuged at 400 x g for 5 minutes at 4°C unless otherwise indicated.

Results were analyzed using FlowJo™ Software (v11). Unstained samples were used to establish the gating strategy, FMO controls were used to define gating boundaries, and single-stained controls were used to correct for background fluorescence. Antibodies and other materials used in this section are listed in Supplementary Table II.

### 4.6 Immunofluorescence staining of HLOs

To generate samples for immunofluorescence staining, organoids were grown and matured after single-cell seeding in Nunc® Lab-Tek® chamber slide^TM^ systems (Sigma Aldrich, #C7057-1PAK) (one 10 μL dome per well). HLOs were fixated for 30 minutes at RT in 4% PFA in PBS (v/v) (Thermo Fisher Scientific, #28908) at the end of the expansion stage (EM Day 5) and the end of the maturation stage (MM Day 10). After washing with PBS, samples were permeabilized for 30 minutes at RT in 0.1% Triton X-100 in PBS (v/v) (Sigma Aldrich, #T8787). After permeabilization, blocking was performed in BSA (BSA in DPBS (3% w/v), Sigma Aldrich #A2153) for 1 hour at RT. Primary antibodies were diluted in 3% BSA, and samples were incubated overnight at 4°C. Samples were washed twice with PBS, and secondary antibodies were diluted in 3% BSA and applied for 2 hours at RT in the dark. For all incubations (permeabilization, fixation, and antibody incubation), Lab-Tek® chambers were placed in a shaking platform at 170 rpm. After three times washing with PBS, HLOs were incubated with DAPI for 30 minutes at RT. Samples were then submerged in mounting medium (Vector Laboratories, #H-1000). Images were taken using a Leica SP8 CLSM confocal immunofluorescent microscope (10x and 20x objective) and analyzed using the ImarisViewer Software (v.11.0.0). Antibodies and other materials used in this section are listed in Supplementary Table III.

### 4.7 Albumin quantification

To assess albumin secretion, 24-hour media was collected at EM Day 5 and MM Day 10 for HLOs and at EM Day 5 and MM Day 14 for LoCs and centrifuged (800 x g, 10 min, 4°C) to remove any cell debris before freezing at -80°C. Albumin concentration was quantified using a commercially available ELISA kit (Thermo Fisher, #EHALB) in 1:100 diluted samples. For HLOs, the amount of albumin in each technical replicate was normalized to ATP production using the 3D-Glo cell viability test (Promega, #G9681). For chips, the amount of albumin was calculated from the continuous media perfusion on the same chip over time, with the final amount of albumin corresponding to the top and the bottom channels combined.

### 4.8 CYP3A4 activity test

CYP3A4 enzyme activity was assessed using the Promega Kit CYP3A4-IPA (Promega, #V9001). CYP3A4 activity of HLOs was assessed at EM Day 5 and MM Day 10. HLOs were seeded in 96-well plates at a density of 5k cells/5 μL Matrigel dome, washed with PBS, and the corresponding media containing the CYP3A4-IPA substrate was added, following the manufacturer’s instructions. Cells were incubated for 18 hours with the CYP3A4-IPA substrate, media was collected, and the luminescence readout corresponding to CYP3A4 enzyme activity was measured, following the manufacturer’s instructions. Luminescence results were normalized to ATP production using the 3D-Glo cell viability test (Promega, #G9681), performed on the same wells from which the supernatant was collected after incubation with the CYP3A4 substrate, following the manufacturer’s instructions.

The chips were disconnected from the flow at MM Day 13, and the top channels were incubated with the corresponding media containing CYP3A4 substrate. The chips were incubated statically for 18 hours, and the luminescence and viability readouts were taken at MM Day 14 in the same way described for the HLOs.

### 4.9 Statistics

Statistical analyses were performed in R (v.4.5.3) using the packages tidyverse (v. 2.0.0), *lme4* (v.2.0-1), and *lmerTest* (v.3.2-1). Raw replicate measurements from three independent donors were analyzed across two experimental stages: expansion and maturation. For each donor and stage, the mean and standard deviation (SD) were calculated from replicate measurements. To assess the effect of differentiation stage on albumin production and CYP3A4 activity while accounting for donor-to-donor variability, a linear mixed-effects model was applied using the *lmer* function. In the model, experimental stage was included as a fixed effect and donor identity was included as a random intercept term (Value ∼ Stage + (1 | Donor)). Statistical significance was determined from the mixed-effects model output.

### 4.10 RNA isolation and sequencing

RNA was isolated from the Matrigel domes using the RNasy MiniKit (#74106, Qiagen). After washing the domes with PBS, 700 μL of lysis buffer was added on top of the domes, and the organoids were mechanically dissociated until no cell clumps were visible. Samples were stored at -80°C until further processing. RNA isolation was performed following the manufacturer’s protocol.

For the LoCs, RNA was isolated using the RNasy MicroKit (#79254, Qiagen). On maturation day 14, the chips were disconnected from the flow, and each channel was washed with 200 μL of PBS. Lysis buffer was added to the channel, and cells were vigorously triturated and collected in an Eppendorf tube. This process was repeated with another 150 μL of lysis buffer until all the cells from the top channel were lysed. Cells were then vortexed and stored at -80°C until further processing. RNA isolation was performed following the manufacturer’s instructions. RNA integrity (RIN > 8) was confirmed using the Agilent RNA ScreenTape (#5067-5576) on the Agilent TapeStation 4200, and the samples were stored at -80°C until sequencing. RNA library preparation and sequencing were performed in two batches (the organoids and the chips) at Novogene (Germany).

### 4.11 Gene expression quantification and quality control

Raw paired-end sequencing reads were obtained in FASTQ format, and quality control was performed using FastQC (v0.12.1) [18]. Transcripts were quantified using Salmon (v1.10.3) in selective-alignment mode (--validateMappings) against a decoy-aware GRCh38 transcriptome index (Ensemble release 86, k = 31, 195 decoys) [19]. Transcript-level abundances were imported into R and summarized to gene-level counts with tximport (v1.32.0, length-scaled TPM) [20].

### 4.12 Differential gene expression analysis

Differentially expressed genes (DEGs) between different conditions and systems were identified using DESeq2 (v1.44.0). These analyses were structured around two main contrasts:

1. *Expansion vs maturation stage*. Prior to filtering, genes were filtered to include only those with at least 300 reads in 50% of samples. The design formula used was: ∼ Donor + Stage.
2. *Organoid vs LoC.* Prior to filtering, genes were filtered to include only those with at least 10 reads in 50% of samples. Principal component 1 (PC1) from a principal component analysis (PCA) of variance-stabilized counts was included as a covariate to adjust for dominant latent variation reflecting inter-donor and technical structure, resulting in the design formula: ∼ PC1 + System.

Benjamini-Hochberg adjustment was applied, and log2 fold changes were shrunk with apeglm (v1.26.1) [21]. DEGs were filtered on having an absolute log2FoldChange ≥ 1 and an adjusted p-value < 0.05.

### 4.13 Pathway analysis

Over-representation analysis was performed separately for the DEGs upregulated and downregulated in each contrast using R package clusterProfiler (v4.12.6). Gene Ontology Biological Process (GO BP) enrichment was assessed with enrichGO using human gene annotations from org.Hs.eg.db (v3.19.1) and Gene Ontology from GO.db (v3.19.1). P-values were adjusted using the Benjamini-Hochberg procedure. GO results were simplified where appropriate (semantic similarity cutoff 0.7) and visualized as dot plots and enrichment maps (enrichplot v1.24.4: emapplot, with pairwise_termsim). All analyses were run in R (version 4.4.1).

### 4.14 Code availability

https://github.com/GRONINGEN-MICROBIOME CENTRE/Chips_vs_Organoids_Tamargo_2026

## Supporting information

Suppementary Material

## Abbreviations

CYP450: cytochrome P450 family
DE: definitive endoderm
DEG: differentially expressed gene
ECM: extracellular matrix
HLO: human liver organoid
hiPSC: human induced pluripotent stem cell
HP: hepatic progenitor
LoC: liver-on-a-chip
OoC: organ-on-a-chip

## Acknowledgements

The authors thank Kate Mc Intyre for editing the manuscript and Iwan J. Hidding for helping with statistical analysis. We used BioRender to generate part of the figures, using a personal BioRender account of one of the authors.

## Data Availability Statement

The data supporting the findings of this study are available from the corresponding author upon reasonable request. The code used for data processing and analysis is publicly available on GitHub at <u>GRONINGEN-MICROBIOME-CENTRE/Chips_vs_Organoids_Tamargo_2026</u>⍰

## Authors contributions

J.F. and S.W. supervised the project. J.A.H. supervised the data analysis and interpretation and contributed to the experimental design and manuscript preparation. I.T.R. conceived the project, designed and conducted the experiments, wrote the manuscript, and prepared the figures. I.T.R. and T.K. analyzed the data. T.K. wrote the Methods section describing the RNA-seq analysis and prepared the corresponding data visualizations. C.v.d.W. helped generate the ELISA data. R.M., G.D.L.W., B.G., and V.E.J.M.P. contributed to optimization experiments performed prior to the generation of the data presented in this study. All authors reviewed the manuscript and approved the final version.

