## Supplementary material for "Comparison of hiPSC-derived hepatic organoids and liver-on-a-chip systems reveal microenvironment-driven maturation": Suppementary Material

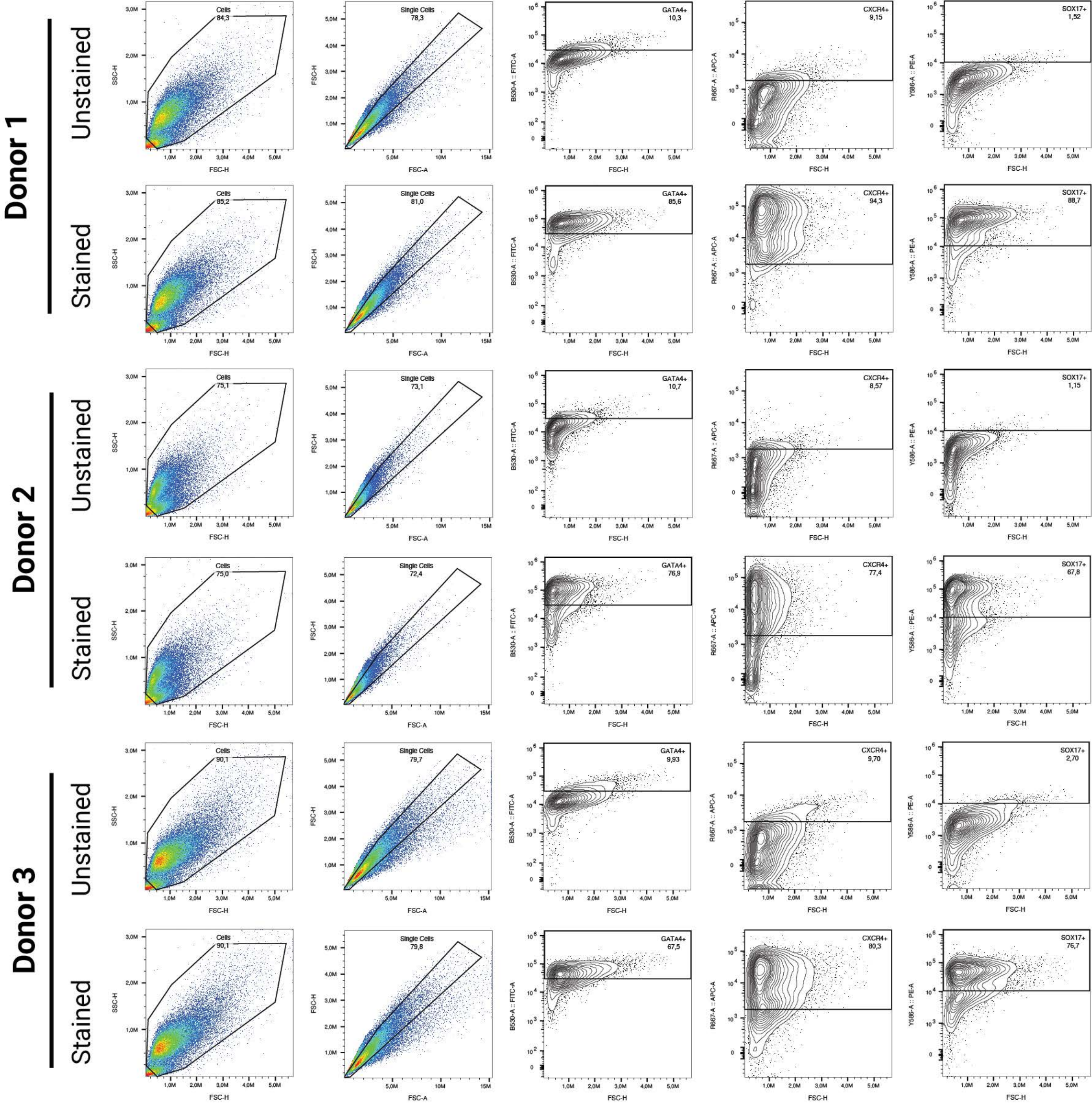

Donor 1
Unstained
Cells 84,3
Single Cells 78,3
GATA4+ 10,3
CXCR4+ 9,15
SOX17+ 1,52
Stained
Cells 85,2
Single Cells 81,0
GATA4+ 85,6
CXCR4+ 94,3
SOX17+ 88,7
Donor 2
Unstained
Cells 75,1
Single Cells 73,1
GATA4+ 10,7
CXCR4+ 8,57
SOX17+ 1,15
Stained
Cells 75,0
Single Cells 72,4
GATA4+ 76,9
CXCR4+ 77,4
SOX17+ 67,8
Donor 3
Unstained
Cells 90,1
Single Cells 79,7
GATA4+ 9,93
CXCR4+ 9,70
SOX17+ 2,70
Stained
Cells 90,1
Single Cells 79,8
GATA4+ 67,5
CXCR4+ 80,3
SOX17+ 76,7
SSC-H
FSC-H
FSC-A
B530-A :: FITC-A
R667-A :: APC-A
Y586-A :: PE-A

**Supplementary Figure 1. Definitive endoderm induction characterization by flow cytometry for three hiPSC donors.** The gating strategy for each donor is selected based on each donor's unstained control.

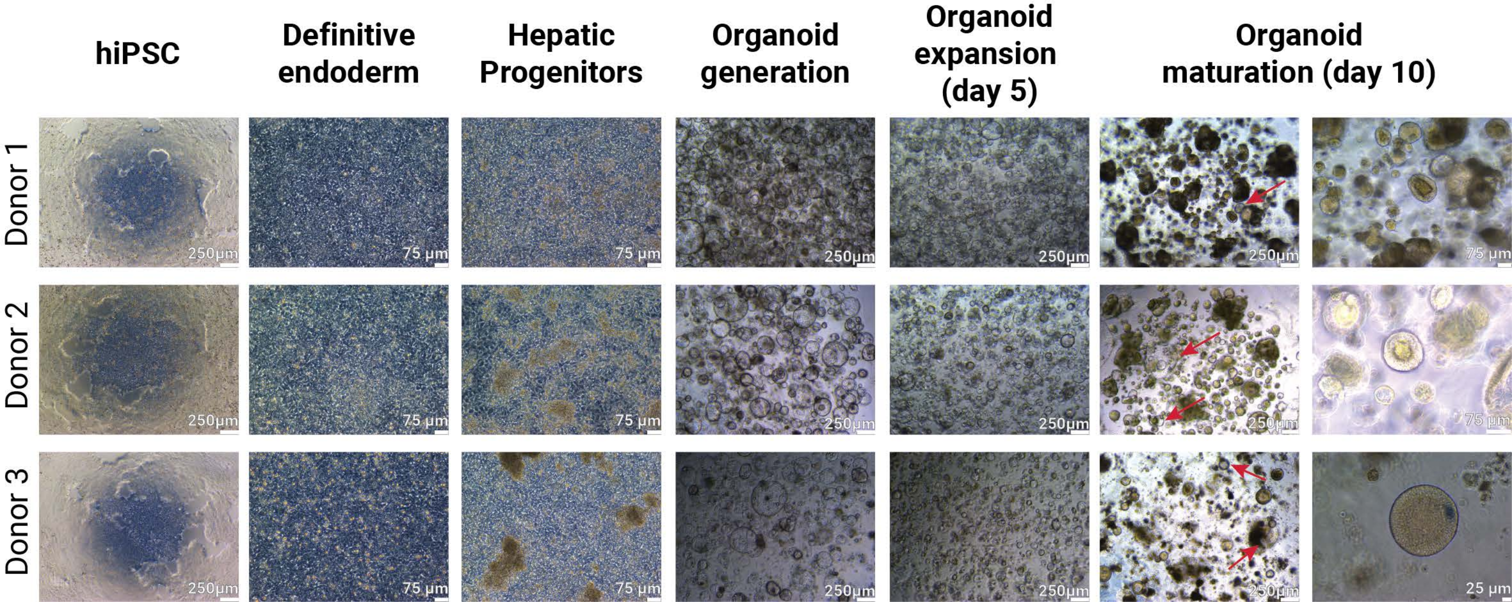
hiPSC
Definitive endoderm
Hepatic Progenitors
Organoid generation
Organoid expansion (day 5)
Organoid maturation (day 10)
Donor 1
Donor 2
Donor 3
250μm
75 μm
25 μm

**Supplementary Figure 2. Bright field pictures of the differentiation protocol from hiPSCs to mature HLOs at the end of each differentiation stage for three hiPSC donors.** Red arrows indicating hollow organoid structures present at the end of the maturation

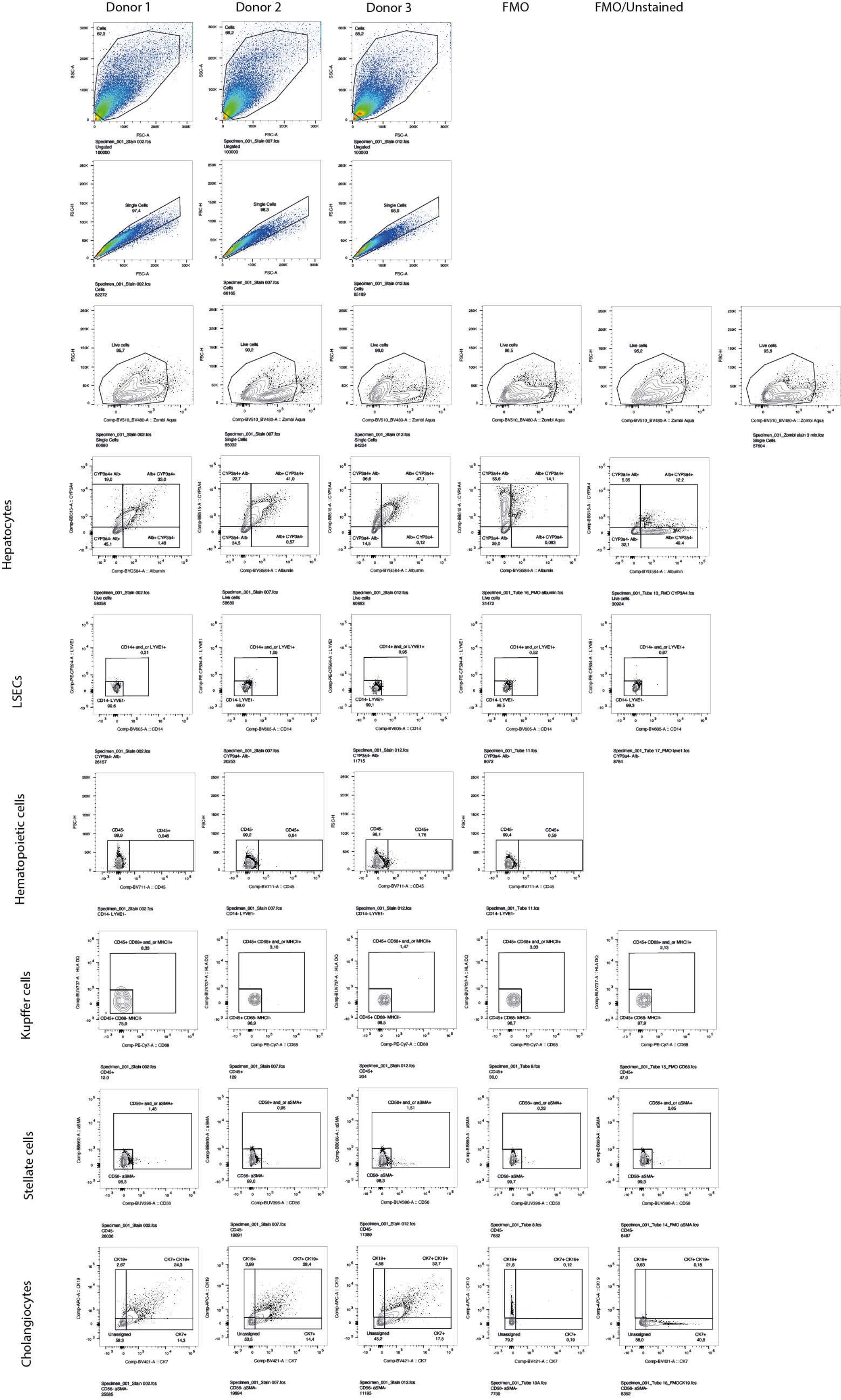

Donor 1
Donor 2
Donor 3
FMO
FMO/Unstained
Hepatocytes
LSECs
Hematopoietic cells
Kupffer cells
Stellate cells
Cholangiocytes

**Supplementary Figure 3. Gating strategy for liver-specific cell type composition in mature HLOs by flow cytometry in three hiPSC donors.** FMO; fluorescence-minus-one, LSECs; liver sinusoidal epithelial cells.

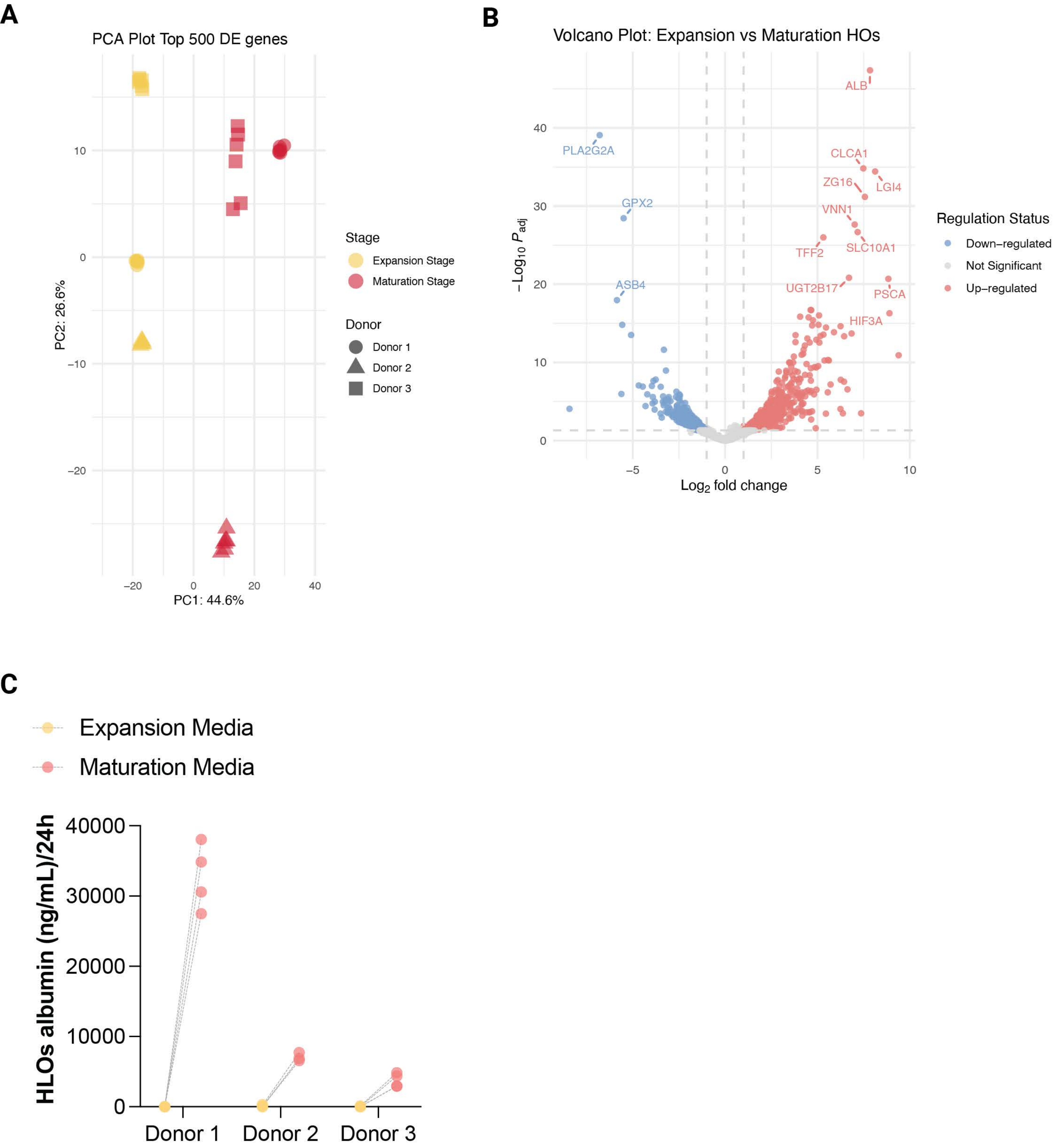
A
PCA Plot Top 500 DE genes
PC1: 44.6%
PC2: 26.6%
Stage
Expansion Stage
Maturation Stage
Donor
Donor 1
Donor 2
Donor 3
B
Volcano Plot: Expansion vs Maturation HOs
$-Log_{10}\ P_{adj}$
$Log_2$ fold change
ALB
PLA2G2A
CLCA1
ZG16
LGI4
GPX2
VNN1
TFF2
SLC10A1
ASB4
UGT2B17
PSCA
HIF3A
Regulation Status
Down–regulated
Not Significant
Up–regulated
C
Expansion Media
Maturation Media
HLOs albumin (ng/mL)/24h
Donor 1
Donor 2
Donor 3

**Supplementary Figure 4. Differential gene expression analysis in HLOs between the end of the expansion and maturation stages for three hiPSC donors and albumin production per donor. (A)** Principal component analysis (PCA) plot for HLOs in the expansion (yellow) and maturation (red) stages. **(B)** Volcano plot displaying the top 10 upregulated and downregulated genes for HLOs in the expansion and maturation stages. N = 6 technical replicates per donor and stage. **(C)** Increase in albumin (ng/mL) secretion from HLOs over time (in the same sample) from the end of the expansion stage to the end of the maturation stage. N = 4 technical replicates per donor and stage

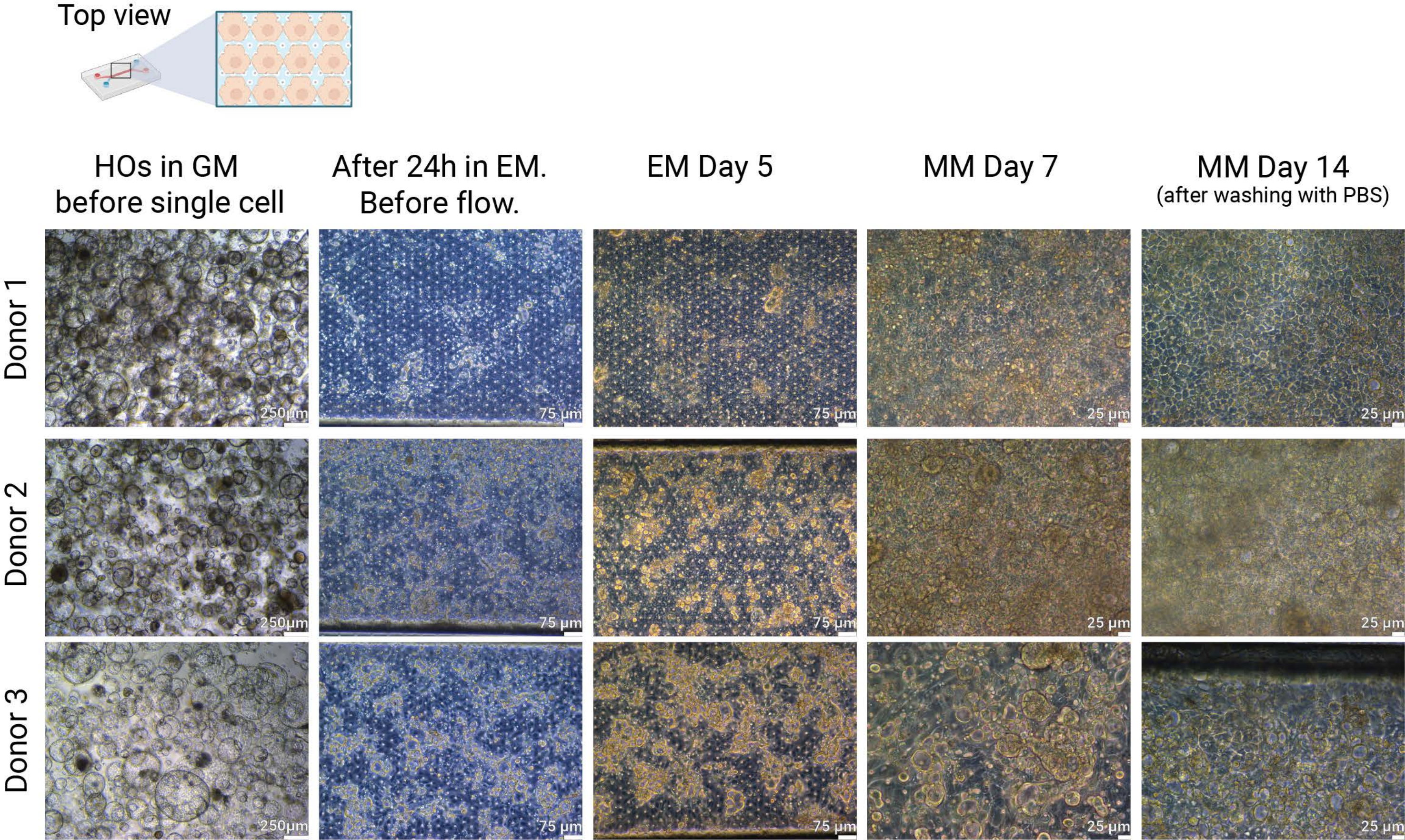

Top view
HOs in GM before single cell
After 24h in EM. Before flow.
EM Day 5
MM Day 7
MM Day 14 (after washing with PBS)
Donor 1
Donor 2
Donor 3
250µm
75 µm
25 µm

**Supplementary Figure 5. Bright field pictures of the differentiation stages from HLOs in generation media (GM) to mature LoCs.** Pictures were taken from a top view (as schematically represented) at four different timepoints throughout the differentiation: 24h after seeding in expansion media (EM), at the end of the expansion stage (EM Day 5), midway in the maturation stage (MM Day 7) and at the end of the maturation stage (MM Day 14) for three hiPSC donors.

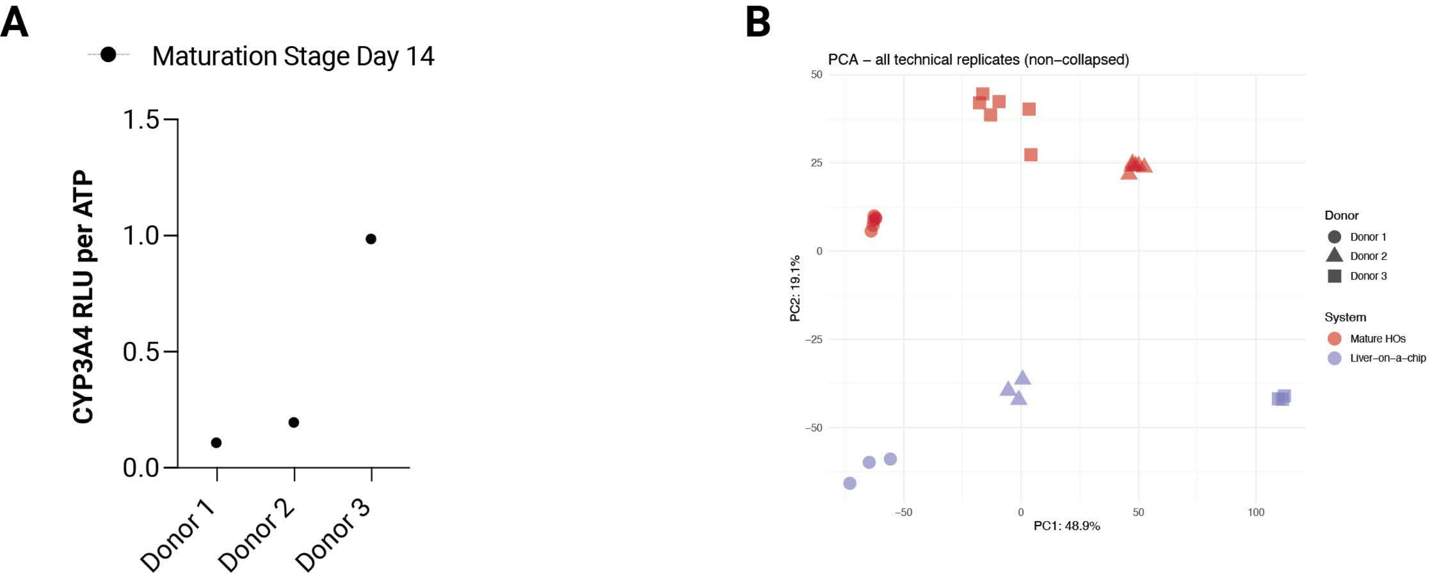
A
Maturation Stage Day 14
CYP3A4 RLU per ATP
1.5
1.0
0.5
0.0
Donor 1
Donor 2
Donor 3
B
PCA – all technical replicates (non–collapsed)
50
25
0
−25
−50
−50
0
50
100
PC1: 48.9%
PC2: 19.1%
Donor
Donor 1
Donor 2
Donor 3
System
Mature HOs
Liver–on–a–chip

**Supplementary Figure 6. CYP3A4 activity on-chip and differential gene expression analysis between mature HLOs and mature LoCs. (A)** CYP3A4 activity measured at the end of the maturation stage (MMD14) in LoCs (n = 1 per donor). Values are corrected for ATP per technical replicate. **(B)** PCA plot for LoCs (purple) and HLOs (red) at the end of the maturation stage for three hiPSC donors. Technical replicates per donor: n = 3 for LoC and n= 6 for HLOs.

**Supplementary Table I.** Composition of media used during the differentiation process including final concentration and product details.

| MEDIA TYPE or COMPONENT | FINAL CONCENTRATION | PRODUCT DETAILS |
|---|---|---|
| Definitive endoderm (DE) day 1 | | |
| RPMI 1640 | | Thermo Scientific Cat# 11875093 |
| L-glutamine | 2mM | Thermo Scientific Cat# 25030081 |
| Pen/Strep | 100U/mL | Thermo Scientific Cat# 15140122 |
| Activin A | 100ng/mL | R&D Cat# 338-AC-010 |
| Wnt3a | 25ng/mL | R&D Cat# 5036-WN/CF |
| Definitive endoderm (DE) day 2 | | |
| RPMI 1640 | | Thermo Scientific Cat# 11875093 |
| L-glutamine | 2mM | Thermo Scientific Cat# 25030081 |
| Pen/Strep | 100U/mL | Thermo Scientific Cat# 15140122 |
| Activin A | 100ng/mL | R&D Cat# 338-AC-010 |
| dFBS | 0.2% (v/v) | Hyclone Cat# SH30070.02 |
| Definitive endoderm (DE) day 3 | | |
| RPMI 1640 | | Thermo Scientific Cat# 11875093 |
| L-glutamine | 2mM | Thermo Scientific Cat# 25030081 |
| Pen/Strep | Pen 100 Units/ml; strep 100 ug/ml | Thermo Scientific Cat# 15140122 |
| Activin A | 100ng/mL | R&D Cat# 338-AC-010 |
| dFBS | 2% (v/v) | Hyclone Cat# SH30070.02 |
| Hepatic progenitor induction (HP) | | |
| RPMI 1640 | | Thermo Scientific Cat# 11875093 |
| B27 | 1x | Thermo Scientific Cat#17504001 |
| BMP4 | 20 ng/mL | R&D Cat#314-BP-CF |
| FGF2 | 10ng/mL | R&D Cat#233-FB/CF |
| DMSO | 0.5% (v/v) | Sigma Aldrich Cat# D2650 |
| Generation Media (GM) | | |
| Adv. DMEM/F12 | | Thermo Scientific Cat# 12634028 |
| GlutaMAX | 1x | Thermo Scientific Cat# 35050061 |
| HEPES | 10mM | Thermo Scientific Cat# 15630080 |

| | | |
|---|---|---|
| N2 | 1x | Thermo Fisher Cat#17502048 |
| B27 | 1x | Thermo Scientific Cat#17504001 |
| BSA | 0.1% | Sigma Aldrich Cat#17502048 |
| N-acetyl L-cysteine | 1.25 mM | Sigma Aldrich Cat#A9165-25G |
| [Leu15]-gastrin I human | 10 nM | Sigma Aldrich Cat#G9145-.1MG |
| Nicotinamide | 10 mM | Sigma Aldrich Cat#N0636-100G |
| A83-01 | 5 uM | R&D Cat#2939/10 |
| Forskolin | 10 uM | R&D Cat#1099/10 |
| CHIR99021 | 3 uM | R&D Cat#4423/10 |
| FGF10 | 50 ng/mL | Peprotech Cat#100-26 |
| HGF | 25 ng/mL | R&D Cat#294-HG/CF |
| Pen/Strep | 50 U/mL | Thermo Scientific Cat# 15140122 |
| Expansion Media (EM) | | |
| Adv. DMEM/F12 | | Thermo Scientific Cat# 12634028 |
| GlutaMAX | 1x | Thermo Scientific Cat# 35050061 |
| HEPES | 10mM | Thermo Scientific Cat# 15630080 |
| N2 | 1x | Thermo Scientific Cat#17502048 |
| B27 | 1x | Thermo Scientific Cat#17504001 |
| BSA | 0.1% | Sigma Aldrich Cat#17502048 |
| N-acetyl L-cysteine | 1.25 mM | Sigma Aldrich Cat#A9165-25G |
| [Leu15]-gastrin I human | 10 nM | Sigma Aldrich Cat#G9145-.1MG |
| Nicotinamide | 10 mM | Sigma Aldrich Cat#N0636-100G |
| A83-01 | 5 uM | R&D Cat#2939/10 |
| Forskolin | 10 uM | R&D Cat#1099/10 |
| CHIR99021 | 3 uM | R&D Cat#4423/10 |
| FGF10 | 50 ng/mL | Peprotech Cat#100-26 |
| HGF | 25 ng/mL | R&D Cat#294-HG/CF |
| BMP7 | 25 ng/mL | R&D Cat#354-BP-010/CF |
| Pen/Strep | 50 U/mL | Thermo Scientific Cat# 15140122 |
| Maturation Media (MM) | | |
| Adv. DMEM/F12 | | Thermo Scientific Cat# 12634028 |
| GlutaMAX | 1x | Thermo Scientific |

| | | |
|---|---|---|
| | | Cat# 35050061 |
| HEPES | 10 mM | Thermo Scientific Cat# 15630080 |
| B27 | 1x | Thermo Scientific Cat#17504001 |
| BSA | 0.1% | Sigma Aldrich Cat#17502048 |
| N-acetyl L-cysteine | 1.25 mM | Sigma Aldrich Cat#A9165-25G |
| [Leu15]-gastrin I human | 10 nM | Sigma Aldrich Cat#G9145-.1MG |
| DAPT | 10 uM | R&D Cat#2634/10 |
| A83-01 | 0.5 uM | R&D Cat#2939/10 |
| Dexamethasone | 3 uM | Sigma Aldrich Cat#D4902 |
| FGF19 | 100 ng/mL | R&D Cat#969-FG-025/CF |
| HGF | 25 ng/mL | R&D Cat#294-HG/CF |
| BMP7 | 25 ng/mL | R&D Cat#354-BP-010/CF |
| Pen/Strep | 50 U/mL | Thermo Scientific Cat# 15140122 |

**Supplementary Table II.** Antibodies for flow cytometry.

| Antigen | Fluorophore | Dilution | Catalogue # | Manufacturer |
|---|---|---|---|---|
| **Definitive endoderm** | | | | |
| CXCR4 | APC | 1:100 | | BD Biosciences |
| GATA4 | Alexa Fluor 488 | 1:100 | | BD Biosciences |
| SOX17 | PE | 1:100 | | BD Biosciences |
| **Mature HOs cell type composition** | | | | |
| CD56 | BUV395 | 1:100 | 563555 | BD Biosciences |
| HLA-DQ | BUV737 | 1:100 | 748562 | BD Biosciences |
| CD14 | BV605 | 1:100 | 564054 | BD Biosciences |
| CD45 | BV711 | 1:100 | 564358 | BD Biosciences |
| LYVE1 | Alexa Fluor 594 | 1:100 | FAB11463T-100UG | R&D |
| CK7 | BV421 | 1:100 | 564709 | BD Biosciences |
| CYP3A4 | FITC | 1:100 | NBP2-50207F | Novus |
| ACTA2 | Percp | 1:100 | IC1420C | R&D |
| CD68 | PE-Cy7 | 1:100 | 565595 | BD Biosciences |
| Albumin | PE | 1:100 | IC1455P | R&D |
| CK19 | APC | 1:100 | ab221255 | abcam |
| HNF4a | Alexa Fluor 750 | 1:100 | IC4605S-100UG | R&D |

**Supplementary Table III.** Primary and secondary antibodies for immunofluorescence stainings.

| Antigen | Dilution | Host/Isotype | Catalogue # | Manufacturer |
|---|---|---|---|---|
| Albumin | 1:200 | Mouse IgG2A monoclonal | MAB1455 | R&D |
| HNF4α | 1:200 | Rabbit IgG monoclonal | C11F12/3113S | Cell Signalling |
| ZO1 | 1:200 | Rabbit IgG polyclonal | 61-7300 | ThermoFisher |
| BCRP1 | 1:100 | Mouse IgG2A monoclonal | ab3380 | abcam |
| **Fluorophore** | **Dilution** | **Species reactivity** | **Catalogue #** | **Manufacturer** |
| Alexa Fluor 488 | 1:250 | Mouse IgG (H+L) | 715-545-150 | Jackson ImmunoResearch |
| Cy3 | 1:250 | Rabbit IgG (H+L) | 711-165-152 | Jackson ImmunoResearch |

**Supplementary Table IV.** Percentages of single positive and double positive cells for all cell types identified in Figure 1E by flow cytometry. FMO; fluorescence-minus-one, LSECs; liver sinusoidal epithelial cells.

| | Cells/Single Cells \| Count | Cells/Single Cells/Live cells \| Count |
|---|---|---|
| Specimen_001_Stain 002.fcs | 60680 | 58058 |
| Specimen_001_Stain 007.fcs | 65032 | 58680 |
| Specimen_001_Stain 012.fcs | 84224 | 80863 |
| Specimen_001_Tube 8.fcs | 33042 | 31171 |
| Specimen_001_Tube 9.fcs | 32794 | 30952 |
| Specimen_001_Tube 10A.fcs | 33555 | 31669 |
| Specimen_001_Tube 10B.fcs | 30838 | 29751 |
| Specimen_001_Tube 11.fcs | 32878 | 31088 |
| Specimen_001_Tube 12.fcs | 32239 | 30482 |
| Specimen_001_Tube 13_FMO CYP3A4.fcs | 32795 | 30924 |
| Specimen_001_Tube 14_FMO aSMA.fcs | 32889 | 31064 |
| Specimen_001_Tube 15_FMO CD68.fcs | 32919 | 31097 |
| Specimen_001_Tube 16_FMO albumin.fcs | 33336 | 31472 |
| Specimen_001_Tube 17_FMO lyve1.fcs | 32641 | 30964 |
| Specimen_001_Tube 18_FMOCK19.fcs | 34173 | 32209 |
| Specimen_001_Tube 19_FMO HNF4A_001.fcs | 32976 | 31224 |
| Specimen_001_UNstained 007.fcs | 6351 | 6006 |
| Specimen_001_UNstained 012.fcs | 8152 | 7986 |
| Specimen_001_Unstained 002.fcs | 5951 | 5665 |
| Specimen_001_Zombi stain 3 mix.fcs | 57604 | 55052 |
| Mean | 35753 | 33819 |
| SD | 19107 | 18071 |

| | % Hepatocytes like | %LSECs like |
|---|---|---|
| Geni002 | 32,962 | 0,138 |
| Geni007 | 41,024 | 0,375 |
| Geni012 | 47,054 | 0,137 |

| Cells/Single Cells/Live cells/Alb+ CYP3a4+ \| Count |
| ---: |
| 19137 |
| 24073 |
| 38049 |
| 18247 |
| 15532 |
| 18677 |
| 10344 |
| 18274 |
| 14264 |
| 3758 |
| 15323 |
| 16634 |
| 4447 |
| 17092 |
| 17867 |
| 17102 |
| 613 |
| 245 |
| 641 |
| 3342 |
| 13683 |
| 9397 |

| %Immune Like like |
| ---: |
| 0,002 |
| 0,007 |
| 0,004 |

| Cells/Single Cells/Live cells/Alb+ CYP3a4+/HNF4a+ Alb+ CYP3a4+ \| Count |
|---:|
| 387 |
| 210 |
| 189 |
| 46 |
| 49 |
| 41 |
| 245 |
| 36 |
| 47 |
| 47 |
| 35 |
| 54 |
| 31 |
| 36 |
| 34 |
| 41 |
| 11 |
| 24 |
| 2 |
| 31 |
| 79,8 |
| 98,7 |

| %Cholangiocytes like |
|---:|
| 10,710 |
| 9,521 |
| 4,532 |

| Cells/Single Cells/Live cells/Alb+ CYP3a4- \| Count | Cells/Single Cells/Live cells/CYP3a4+ Alb- \| Count |
|---|---|
| 857 | 11025 |
| 336 | 13332 |
| 98 | 29581 |
| 420 | 4095 |
| 167 | 6487 |
| 428 | 4201 |
| 188 | 7788 |
| 482 | 3847 |
| 222 | 7116 |
| 15291 | 1655 |
| 262 | 6337 |
| 396 | 4783 |
| 26 | 17494 |
| 569 | 4097 |
| 275 | 5066 |
| 400 | 4626 |
| 4 | 780 |
| 1 | 440 |
| 6 | 794 |
| 34 | 6025 |
| 1023 | 6978 |
| 3366 | 6791 |

| %Stellate like | Unassigned | |
|---|---|---|
| 0,641 | | 25,709 |
| 0,320 | | 17,969 |
| 0,213 | | 6,256 |

| Cells/Single Cells/Live cells/CYP3a4- Alb- \| Count |
| ---: |
| 26157 |
| 20253 |
| 11715 |
| 7987 |
| 8297 |
| 7879 |
| 11084 |
| 8072 |
| 8344 |
| 9921 |
| 8606 |
| 8922 |
| 9132 |
| 8784 |
| 8545 |
| 8646 |
| 4583 |
| 7287 |
| 4192 |
| 45412 |
| 11691 |
| 9360 |

| unassigned + single positives |
| ---: |
| 31241,000 |
| 28176,000 |
| 37520,000 |

| Cells/Single Cells/Live cells/CYP3a4- Alb-/CD14+ and_or LYVE1+ \| Count |
| ---: |
| 80 |
| 220 |
| 111 |
| 56 |
| 45 |
| 50 |
| 68 |
| 42 |
| 52 |
| 68 |
| 58 |
| 48 |
| 49 |
| 59 |
| 45 |
| 95 |
| 28 |
| 11 |
| 12 |
| 144 |
| 67 |
| 47,6 |
| 53,810 |
| 48,016 |
| 46,399 |

| Cells/Single Cells/Live cells/CYP3a4- Alb-/CD14- LYVE1- \| Coun |
|---|
| 26061 |
| 20042 |
| 11609 |
| 7933 |
| 8245 |
| 7827 |
| 11014 |
| 8034 |
| 8287 |
| 9841 |
| 8547 |
| 8867 |
| 9076 |
| 8719 |
| 8497 |
| 8554 |
| 4556 |
| 7275 |
| 4178 |
| 45275 |
| 11622 |
| 9331 |

Cells/Single Cells/Live cells/CYP3a4- Alb-/CD14- LYVE1-/CD45+ | Coun

12
129
204
44
30
44
61
47
38
53
58
47
42
40
40
44
14
25
1
97
53,5
45,2

| Cells/Single Cells/Live cells/CYP3a4- Alb-/CD14- LYVE1-/CD45+/CD45+ CD68+ and_or MHCII+ \| Cour |
|---:|
| 1 |
| 4 |
| 3 |
| 0 |
| 1 |
| 1 |
| 5 |
| 1 |
| 0 |
| 4 |
| 3 |
| 1 |
| 3 |
| 1 |
| 0 |
| 1 |
| 0 |
| 0 |
| 0 |
| 1 |
| 1,5 |
| 1,57 |

| Cells/Single Cells/Live cells/CYP3a4- Alb-/CD14- LYVE1-/CD45+/CD45+ CD68- MHCII- \| Cour |
|---:|
| 9 |
| 125 |
| 201 |
| 44 |
| 29 |
| 43 |
| 55 |
| 46 |
| 38 |
| 49 |
| 54 |
| 46 |
| 39 |
| 39 |
| 40 |
| 43 |
| 14 |
| 25 |
| 1 |
| 96 |
| 51,8 |
| 44,5 |

| Cells/Single Cells/Live cells/CYP3a4- Alb-/CD14- LYVE1-/CD45- \| Coun |
|---|
| 26036 |
| 19891 |
| 11389 |
| 7882 |
| 8211 |
| 7778 |
| 10944 |
| 7982 |
| 8246 |
| 9773 |
| 8487 |
| 8813 |
| 9029 |
| 8676 |
| 8454 |
| 8507 |
| 4499 |
| 7202 |
| 4094 |
| 44890 |
| 11539 |
| 9267 |

| Cells/Single Cells/Live cells/CYP3a4- Alb-/CD14- LYVE1-/CD45-/CD56+ and_or aSMA+ \| Coun |
|---:|
| 372 |
| 188 |
| 172 |
| 26 |
| 58 |
| 38 |
| 101 |
| 45 |
| 59 |
| 104 |
| 55 |
| 52 |
| 78 |
| 58 |
| 98 |
| 46 |
| 57 |
| 51 |
| 20 |
| 321 |
| 100 |
| 95,1 |

| Cells/Single Cells/Live cells/CYP3a4- Alb-/CD14- LYVE1-/CD45-/CD56- aSMA- \| Cour |
| ---: |
| 25585 |
| 19694 |
| 11195 |
| 7856 |
| 8151 |
| 7739 |
| 10835 |
| 7936 |
| 8185 |
| 9651 |
| 8431 |
| 8757 |
| 8946 |
| 8615 |
| 8352 |
| 8459 |
| 4437 |
| 7140 |
| 4070 |
| 44531 |
| 11428 |
| 9170 |

| Cells/Single Cells/Live cells/CYP3a4- Alb-/CD14- LYVE1-/CD45-/CD56- aSMA-/CK7+ \| Cour |
|---|
| 3658 |
| 2829 |
| 1954 |
| 1715 |
| 1975 |
| 15 |
| 806 |
| 1685 |
| 1685 |
| 2018 |
| 1768 |
| 2412 |
| 1685 |
| 1581 |
| 3406 |
| 2055 |
| 9 |
| 4 |
| 20 |
| 76 |
| 1568 |
| 1110 |

| Cells/Single Cells/Live cells/CYP3a4- Alb-/CD14- LYVE1-/CD45-/CD56- aSMA-/CK19+ \| Cour |
| ---: |
| 683 |
| 786 |
| 513 |
| 94 |
| 94 |
| 1688 |
| 673 |
| 120 |
| 139 |
| 193 |
| 144 |
| 84 |
| 147 |
| 153 |
| 53 |
| 103 |
| 53 |
| 12 |
| 1 |
| 148 |
| 294 |
| 405 |

| Cells/Single Cells/Live cells/CYP3a4- Alb-/CD14- LYVE1-/CD45-/CD56- aSMA-/CK7+ CK19+ \| Cour |
|---:|
| 6218 |
| 5587 |
| 3665 |
| 2050 |
| 1603 |
| 9 |
| 2812 |
| 1621 |
| 1536 |
| 2895 |
| 1727 |
| 1726 |
| 2003 |
| 1698 |
| 15 |
| 1710 |
| 0 |
| 0 |
| 0 |
| 2 |
| 1844 |
| 1764 |

| Cells/Single Cells/Live cells/CYP3a4- Alb-/CD14- LYVE1-/CD45-/CD56- aSMA-/Unassigned \| Count |
|---:|
| 14926 |
| 10544 |
| 5059 |
| 3965 |
| 4433 |
| 6126 |
| 6571 |
| 4494 |
| 4811 |
| 4519 |
| 4772 |
| 4500 |
| 5093 |
| 5150 |
| 4845 |
| 4560 |
| 4383 |
| 7124 |
| 4047 |
| 44329 |
| 7713 |
| 9005 |
